# Effects of altered cellular ultrastructure on energy metabolism in diabetic cardiomyopathy – an in-silico study

**DOI:** 10.1101/2022.05.22.492785

**Authors:** Shouryadipta Ghosh, Giovanni Guglielmi, Ioannis Orfanidis, Fabian Spill, Anthony Hickey, Eric Hanssen, Vijay Rajagopal

## Abstract

Diabetic cardiomyopathy is a leading cause of heart failure in diabetes. At the cellular level, diabetic cardiomyopathy leads to altered mitochondrial energy metabolism and cardiomyocyte ultrastructure. We combined electron microscopy and computational modelling to understand the impact of diabetes induced ultrastructural changes on cardiac bioenergetics.

We collected transverse micrographs of multiple control and type I diabetic rat cardiomyocytes using electron microscopy. Micrographs were converted to finite element meshes, and bioenergetics was simulated over them using a biophysical model. The simulations also incorporated depressed mitochondrial capacity for oxidative phosphorylation and creatine kinase reactions to simulate diabetes induced mitochondrial dysfunction.

Analysis of micrographs revealed a 14% decline in mitochondrial area fraction in diabetic cardiomyocytes, and an irregular arrangement of mitochondria and myofibrils. Simulations predicted that this irregular arrangement, coupled with depressed activity of mitochondrial creatine kinase enzymes, leads to large spatial variation in ADP/ATP profile of diabetic cardiomyocytes. However, when spatially averaged, myofibrillar ADP/ATP ratios of a cardiomyocyte do not change with diabetes. Instead, average concentration of inorganic phosphate rises by 40% due to lower mitochondrial area fraction and dysfunction in oxidative phosphorylation. These simulations indicate that a disorganized cellular ultrastructure negatively impacts metabolite transport in diabetic cardiomyopathy.

## Introduction

Type I diabetes (T1D) accounts for 5 – 10% of all diabetes cases every year (1, 2). T1D can lead to a variety of cardiovascular complications eventually resulting in heart failure. T1D cardiomyopathy is one of the T1D induced disease processes, commonly associated with left ventricular diastolic dysfunction with normal ejection fraction (3). It exhibits many common metabolic conditions accompanying heart failure, e.g., increased synthesis of reactive oxygen species (ROS) (4, 5), impaired mitochondrial oxidative phosphorylation (OXPHOS) (5-7) and decrease in both cellular reserve of phosphocreatine (PCr) and activity of mitochondrial creatine kinase (mtCK) enzymes regulating PCr levels (8, 9). T1D cardiomyopathy is also accompanied by alterations in the ultrastructure of cardiomyocytes. Cardiomyocytes are densely packed with three-dimensional mitochondrial networks (10, 11) and columns of myofibrils that traverse parallelly across the length of cells. The mitochondrial networks are formed by close contact between the outer membrane of two adjacent mitochondria (10-13). In T1D cardiomyopathy this columnar ultrastructure is altered with changes both in morphology and organization of mitochondria and myofibrils (6, 14, 15). Similar changes are also observed in several other pathological conditions of the heart (16-18). However, the functional role or consequence of these ultrastructural alterations is unclear. In this study, we use type 1 diabetic (T1D) cardiomyopathy as a model disease state to understand how and to what extent the accompanying changes in cardiomyocyte ultrastructure can influence cellular energy metabolism.

The majority of previous studies point towards increased mitochondrial fission as the key mechanism underlying the change in mitochondria organisation and morphology in T1D cardiomyopathy. Increased mitochondrial fission in T1D cardiomyopathy is characterized by smaller mitochondria with higher numeric density (19-21). Our recent study on streptozotocin (STZ) induced T1D in Sprague Dawley (SD) rats further showed that fragmented individual mitochondria cluster together to form mitochondrial clusters of varying shapes and sizes (21). Several studies also report mitochondrial proliferation in the form of higher mitochondrial volume fraction in T1D cardiomyocytes (6, 15, 22). However, these reports conflict with few other studies where mitochondrial content is found to be either unchanged (23) or decreased (24, 25) with T1D. A potential reason behind these conflicting results can be the use of small regions of interest (ROI) on electron microscopy (EM) images. EM micrographs of small ROIs provide great insights at a local level but lacks insights on how mitochondrial organization is affected across the cell. In addition to changes in mitochondrial organization, T1D can also disrupt the organization of other organelles such as myofibrils, T-tubules and sarcoplasmic reticulum (SR) (24, 26).

In our unpublished work on STZ induced T1D in SD rats, substantial changes in cardiomyocyte ultrastructure were only observed 8-9 weeks after injection of STZ. In another relevant study on Alloxan induced T1D in SD rats, Thomson et al. (26) observed that only ∼15% of all cardiomyocytes in the left ventricle undergo structural disorganization after 6 weeks of diabetes. The percentage of disorganized cardiomyocyte increases up to 60% after 26 weeks. In contrast, 4 weeks post STZ injection is sufficient to observe significant changes in cardiac mitochondrial metabolism, e.g., decline in mitochondrial state III oxygen consumption (7) and activity of mtCK enzymes (9, 27). Are the subsequent ultrastructural changes adaptive in nature or do they compound the negative consequences of disrupted mitochondrial metabolism?

A previous study by Shen et al. (14) on OVE26 mice suggests mitochondrial proliferation to be an adaptive response to mitochondrial dysfunction such as increased ROS production. However, two other works on insulin resistant mice indicate that higher mitochondrial volume fraction and DNA content might not be able to compensate for a decrease in respiratory capacity of individual mitochondria (28, 29). In our recent in-silico analysis of energy metabolism in control cardiomyocytes, we found that CK mediated rapid phospho-transfer can maintain near uniform ATP and ADP levels across a cell cross section, despite a non-uniform arrangement of mitochondrial and myofibrillar columns (30). Since CK enzyme activity is reported to be substantially lower in T1D cardiomyocytes (9, 27), disorganization of organelles might negatively impact the cellular mechano-energetic landscape.

The primary aim of the current study was to investigate how and to what extent various ultrastructural alterations accompanying T1D cardiomyopathy can influence the energy metabolism of cardiomyocytes. We utilized a combination of EM imaging and in-silico modelling that allowed us to decouple the interactions between ultrastructural alterations and alterations in mitochondrial metabolism and examine their cumulative effects on cardiac bioenergetics. We first collected and analysed EM images of entire cross sections of cardiomyocytes from control and STZ induced T1D rats. Next, we used our finite element (FE) model of cardiac bioenergetics (30, 31) to simulate both control and diabetic bioenergetics over spatially realistic FE meshes derived from these cross sectional EM images. The simulation predictions revealed that ultrastructural changes such as lower mitochondrial fraction and irregular mitochondrial arrangement further compound the disruptive effects of preceding metabolic dysfunction such as impaired OXPHOS capacity and lower mtCK enzyme activity. The following sections present the image analysis, formulation of the FE models and the subsequent simulation predictions. Our results support a hypothesis that cardiomyocyte ultrastructural changes found in T1D negatively impact cardiac bioenergetics.

## Methods

### Tissue sample preparation and transmission electron microscopy

The animal procedures in this study followed the guidelines approved by the University of Auckland Animal Ethics Committee (for animal procedures conducted in Auckland, Application No. R826). Six SD rats were randomly distributed into a control and a T1D group and raised in two separate cages. After six weeks, T1D was induced in three six-week-old male SD rats by injecting them with a single dose of STZ (55 mg/kg body wt) in saline medium. The same volume of saline without STZ was injected into the three control animals of same age. The T1D animals were euthanized 9 weeks after the injections, while the control animals were euthanized 7 – 9 weeks after the injections. Following sacrifice, hearts were excised, and 400-600 μm side cubes of tissue from the ventricular mid-wall of each control and diabetic hearts were chemically fixed (2.5% glutaraldehyde, 2% paraformaldehyde, and 50 mM CaCl2 in 0.15 M sodium cacodylate buffer) (32) and processed for standard transmission electron microscopy (TEM). Ultrathin sections of 90 nm thickness were subsequently cut from epoxy resin blocks using a diamond knife and utilized for TEM imaging.

A Tecnai Spirit TEM operated at 120kV was used to acquire the majority of the transverse-view two dimensional micrographs with pixel size varying from 5.6 nm to 13.5 nm. A few sections were also imaged with a smaller pixel size of 2.3 nm using a Tecnai F30 TEM operated at 200 kV. Ultimately, a total of 21 diabetic cardiomyocyte cross-sections and 19 control cardiomyocyte cross-sections (Methods S1) were selected from the collected transverse-view images (a minimum of 5 cross-sections per animal). All the cross section images were acquired away from the cell nucleus (33).

### Image segmentation and analysis of cellular ultrastructure

Trainable Weka Segmentation plugin of ImageJ (34), an open-source EM image processing software package was utilized to roughly segment each cell cross-section into four organelle based regions – (i) mitochondria, (ii) myofibrils, (iii) t-tubules and intracellular vacuoles and (iv) glycogen particles. The rough segmentations were further manually corrected using selection tools available in ImageJ. Fig. 1A and B represent final segmentations of two typical cross sections of control and diabetic cardiomyocytes used in the study. The segmented transverse sections were first analysed for area fractions of the different organelles. E.g., area fraction of mitochondria in each cell cross-section was calculated as a ratio of total number of pixels marked as mitochondria vs. total number of pixels within the cell cross-section. This process was repeated for all the four organelle regions (Fig. 1C). The ratio of mitochondrial area fraction against total mitochondrial and myofibrillar area fraction was defined as mito/myo_global_ of each section

**Figure 1:**
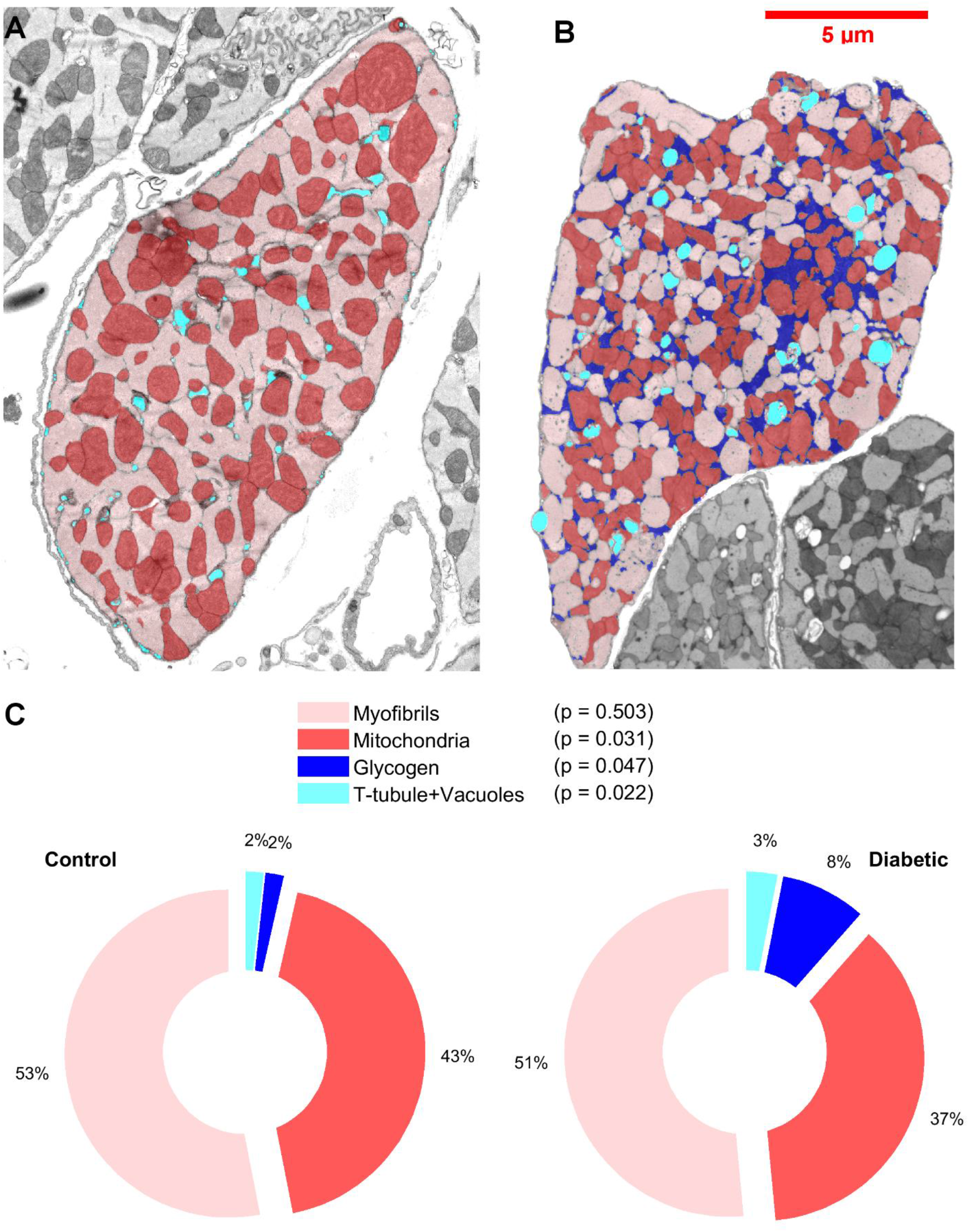
Segmented organelle areas in cardiomyocyte cross-sections and their area fractions. **A**. Segmentation of organelle areas in a representative cardiomyocyte cross-section from TEM micrograph of a control animal. Segmentations are overlaid over original transverse micrograph. **B**. Segmentation of organelle areas in a representative cross-section from a diabetic animal. **C**. Pie charts showing average organelle area fractions in the control and diabetic group. Both groups consist of n = 3 animals, with at least 5 cross sections analysed for each animal. The p-values correspond to the null hypothesis that organelle fractions do not change in diabetes.

Following the calculation of area fractions, distribution of ATP producing sites (mitochondria) with respect to the ATP consuming sites (myofibrils) was analysed for each cell cross-section. In this analysis, a square sampling window of fixed size was assumed to be centred at every pixel marked as myofibril (Fig. S1A). Each sampling window was analysed for the ratio of total mitochondrial pixels against total mitochondrial and myofibrillar pixels present within the window. This ratio was termed as the localized area density of mitochondria (mito/myo_local_) for a given myofibrillar pixel. The side length of the square sampling window was 1.6 μm – which is twice the average diameter of cardiac mitochondria. A further justification of this choice of sampling window length can be found in our previous work (30). Fig. S1A and B maps the resulting mito/myo_local_ distribution in the representative cross sections shown in Fig. 1A and B. The inset in Fig. S1C further shows the histogram representation of these two mito/myo_local_ distributions. To quantify the spatial heterogeneity in mitochondrial-myofibrillar arrangement of a given cross section, median absolute deviation (MAD) of its mito/myo_local_ distribution was calculated. MAD is a robust measure of variability with little sensitivity to presence of outliers.

### Computational modelling of control cardiomyocytes

Our previous publication (31) provides complete mathematical details of the computational model of control cardiomyocytes used in this study, along with the validation of the model predictions with respect to four experimental datasets. The Fortran source code of the model is available at our GitHub repository - https://github.com/CellSMB/cardiac_bioenergetics/tree/V-2.0. The present section provides a brief outline of the model to assist the readers with interpretation of the simulation results. Fig. 2 is a schematic diagram of the different state variables and reaction flux terms that are used in this biophysics based model. Table S1 provides expanded names of the state variables and reaction fluxes used in Fig. 2.

**Figure 2:**
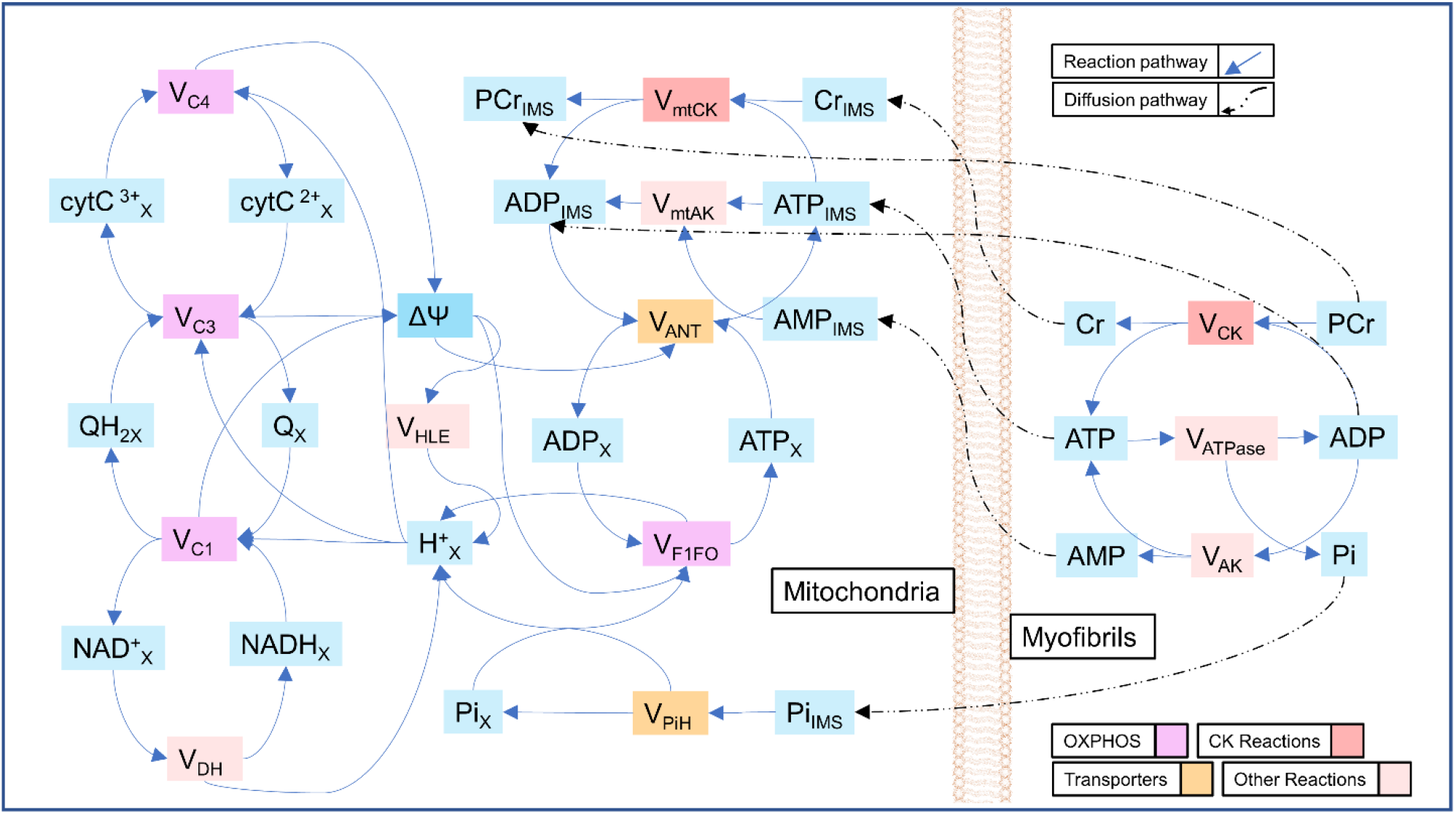
Schematic diagram of the biophysical model of cardiac bioenergetics. The rectangular boxes coloured in cyan represent the state variables of the model. The state variables are molar concentrations of different metabolites present in myofibrils, mitochondrial IMS (suffix - _IMS_) and mitochondrial matrix (suffix - _X_), except for ψ which is mitochondrial membrane potential. All other boxes in the figure represent reaction fluxes involving the state variables. Metabolites present in the myofibrils and mitochondrial IMS also partake in diffusive fluxes across the mitochondria-myofibril interface. Table S1 provides expanded names of the state variables and reaction fluxes used in this figure.

The computational models were based on the TEM micrographs used in the image analysis. The segmented images were first converted into FE meshes using an open source tool called Triangle (35). A previously validated model of OXPHOS in isolated mitochondria (36) was simulated at each mitochondrial node, incorporating differential algebraic equations representing 11 reactions with 13 metabolites (Fig. 2). Key reaction fluxes modelled include: (i) generation of mitochondrial membrane potential and electron transfer through Complex I, III and IV; (ii) generation of ATP at F1-F0 ATP synthase and; (iii) exchange of ATP and ADP through the adenine nucleotide translocases (ANT). However, the isolated mitochondrial model lacked description of reactions occurring in the mitochondrial intermembrane space (IMS) and subsequent diffusion of metabolites through the mitochondrial outer membrane to myofibrils. Therefore, equations were also introduced at each mitochondrial node to simulate key IMS reactions. These include synthesis of PCr catalysed by mtCK, as well as, diffusion of ATP, ADP, Adenosine monophosphate (AMP), PCr, creatine (Cr) and Pi (Fig. 2). The IMS and matrix reactions were modelled as two reaction compartments continuously distributed across the mitochondrial regions. Fluxes of transport reactions such as ANT and phosphate–hydrogen cotransporter mathematically connected both compartments. All reactions fluxes used in the differential equations were scaled by the volume fraction of the respective compartment (VF_IMS_=0.1, VF_MATRIX_=0.9) to maintain conservation of mass.

Similar to the mitochondrial FE nodes, total 5 reactions with 8 metabolites were modelled in each myofibrillar FE node (Fig. 2). Out of these metabolites, concentrations of ATP, ADP and Pi were used to calculate the reaction rate of ATP hydrolysis (**V**_**ATPase**_) in each myofibril nodes (Fig. S2).

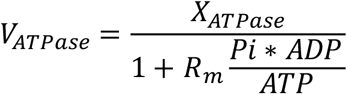

Here R_m_ is a constant of mass-action ratio, while X_ATPase_ is a model input that can be varied to simulate steady state ATP hydrolysis at various workloads. ATP, ADP and Pi were also modelled to diffuse through the myofibrillar nodes towards the mitochondrial nodes. Other key myofibrillar reactions included in the model were regeneration of ATP by myofibrillar creatine kinase (CK) and buffering of ADP/ATP level by myofibrillar adenylate kinase (AK). The reaction rate of CK in myofibrillar nodes was balanced with that of mtCK in mitochondrial nodes through diffusion of PCr and Cr. Similarly, myofibrillar AK reaction rate was balanced with that of mitochondrial AK (mtAK) through diffusion of adenosine monophosphate (AMP). Fig. S2 provides the detailed mathematical equation of few key state variables and reaction fluxes for readers to appreciate the biophysical mechanisms modelled through these equations. The values of the diffusion constants of all diffusing species are available in Table S2.

Glycogen particles usually do not contain any enzymes capable of catalysing reactions involving the metabolites such as ATP and ADP. Therefore, no reaction fluxes were calculated in the glycogen FE nodes, although ATP, ADP, PCr, Cr and Pi were assumed to be diffusing through these nodes with the same diffusivities as those of myofibrillar nodes. It was further assumed that none of the metabolites considered in the model are present in the t-tubules and vacuoles. These regions were marked as holes in the FE mesh. The resulting bioenergetics FE model of a TEM micrograph was simulated using OpenCMISS, an opensource FE modelling software (37).

### Computational modelling of cardiomyocytes with type I diabetes

As discussed earlier, STZ induced T1D cardiomyopathy in SD rats is first accompanied by dysfunction in ATP synthesis machinery of mitochondria within four weeks of STZ injection, followed by alterations in intracellular ultrastructure after another 4 – 5 weeks. Some of the key metabolic changes include: i) decrease in enzymatic activity of several mitochondrial complexes, including complex I (7, 23) and F1-F0 ATP synthase (38, 39), as well as, mtCK present in IMS (9, 27, 40); (ii) elevated level of mitochondrial uncoupling and proton leak (41, 42); (iii) finally, decrease in the level of mitochondrial ATP synthesis (6, 7, 23), membrane potential (43, 44) and O_2_ consumption (42, 45). To understand how ultrastructural alterations interplay with mitochondrial dysfunction, diabetic cardiomyocytes were simulated in two steps. First, diabetic mitochondrial dysfunction was simulated with control cellular ultrastructure (Set CD). Simulations set CD consisted of total 19 simulations based on the 19 control cross sections used for image analysis. Next, diabetic mitochondrial dysfunction was also simulated with FE meshes derived from diabetic TEM images (Set DD). Set DD consisted of total 21 simulations. Results from both simulations set CD and DD were compared with those derived from simulations of control cellular ultrastructure with control mitochondrial metabolism (Set CC, 19 simulations).

Mitochondrial dysfunction in the step CD and DD were simulated by modifying the values of four key parameters present in the mitochondrial model. These parameters are: (i) complex I enzyme activity (X_C1_); (ii) F1-F0 enzyme activity (X_F1_); (iii) proton leak activity (X_HLE_); and (iv) maximal mtCK reaction rates in forward and backward direction (V_1_ and V-_1_). For example, several experimental studies indicate that mtCK enzyme activity (IU/mg myocardial protein) can be decreased by a margin of 35% – 50% in cardiomyocytes after 8 weeks of STZ induced diabetes (9, 27, 40). Based on these studies, both V_1_ and V-_1_ were decreased by a margin of 50% to simulate the diabetic mtCK reactions. Similarly, Pham et al. (5) reported a 35% decrease in mitochondrial ATP synthesis rate, 24% decrease in O_2_ consumption rate and unchanged membrane potential during state III respiration in cardiac tissue homogenates from 8 week STZ induced T1D SD rat hearts. These changes were reproduced in the model by finding values of X_C1_, X_F1_ and X_HLE_ that lead to changes in mitochondrial ATP synthesis rate, O_2_ consumption rate and membrane potential equivalent to that observed in diabetic tissue homogenates. Lsqnonlin, a nonlinear data fitting function in MATLAB (46) was used to estimate these parameters. Table 1 presents the control values of V_1_, V-_1,_ X_C1_, X_F1_ and X_HLE_ alongside the diabetic values of these parameters.

**Table 1:** Control and diabetic values of key OXPHOS parameters altered in the mitochondrial energy metabolism model

| Parameter | Control value | Diabetic value |
| --- | --- | --- |
| $X_{F1}$ ($F1-F0$ activity) | 150.93 | 0.0410 |
| $X_{C1}$ (Complex I activity) | 0.36923 | 0.1063 |
| $X_{HLE}$ (Proton leak activity) | 250 | 437.8554 |

### Statistical Analysis

A linear mixed model (LMM) (47) was used to test if a given variable, e.g. mitochondrial area fraction, shows any difference across the different conditions. For each LMM, the random effect indicates the multiple cross-sections obtained for each rat. The fixed effects of each model are the rat’s conditions, such as control and diabetes for image analysis or CC, CD and DD for simulations results. The parameters of LMM were estimated using the restricted maximum likelihood procedure. The marginal contribution of each condition was tested using the t-statistics.

Correlation between two measurements was quantified by Pearson correlation coefficient, and t-statistics was employed to test if the correlation differs from the null value. The supplementary methods (Methods S1) provides a detailed discussion of the statistical tools used to analyse the model predictions and results from image analysis.

## Results

### Type I diabetes changes area fractions and organization of intracellular organelles

The pie charts in Fig. 1C presents the organelle area fractions averaged over the three animals in both control and diabetic group. It is evident from Fig. 1C that control animals contain a higher fraction of mitochondria (43% in control vs 37% in T1D), while the diabetic animals contain a higher fraction of glycogen particles (2% in control vs 9% in T1D). The average area fraction of myofibrils did not change significantly in diabetic animals. The boxplots in Fig. S1D provides the MADs of the mito/myo_local_ distributions, averaged over control and diabetic animals. The average MAD of mito/myo_local_ distributions increased mildly (0.083 vs. 0.094) in the diabetic group. The higher MAD implies that arrangement of mitochondria and myofibrils is more non-uniform in diabetic cross-sections compared to control cross-sections.

### Myofibrillar ATP Metabolism is altered in Type I diabetes

Fig. 3 presents colour spectrum maps of a few key bioenergetic parameters which regulate cross bridge cycling in myofibrils. These include results from three different simulations (CC, CD and DD) based on the two representative cross sections previously shown in Fig. 1A an B. The same high value of X_ATPase_ (X_ATPase_ = 0.01), indicating a high Ca2+ induced activity of the actomyosin complexes, was used as a model input for the three simulations. This helped in evaluating cardiac bioenergetics independent of alterations in Ca2+ dynamics.

**Figure 3:**
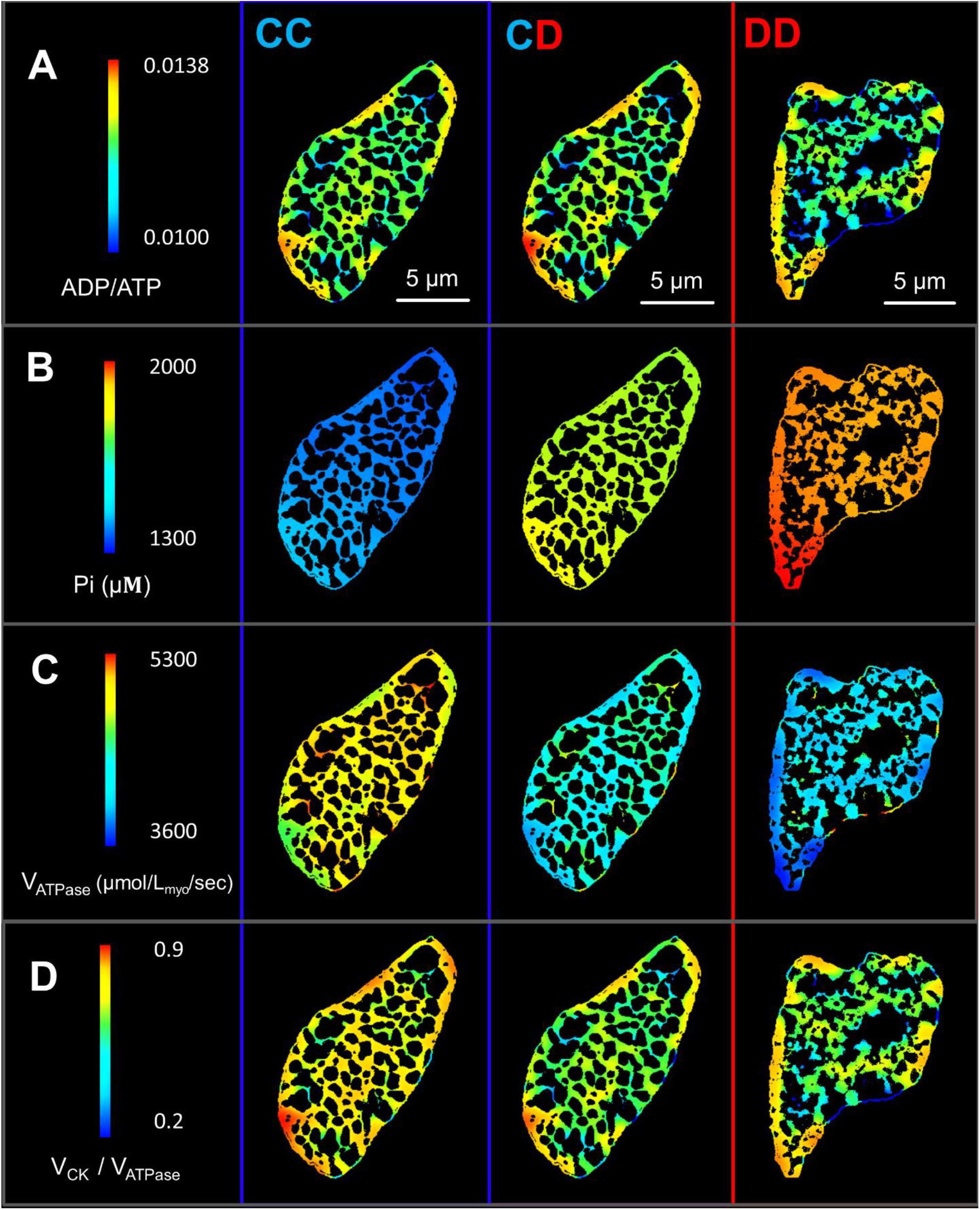
Model predicted spatial distributions of key metabolic parameters in the myofibrillar regions of representative control and diabetic cross sections. Each column of the figure presents results from three distinct simulation sets – control mitochondrial metabolism with control ultrastructure (Set CC – column 1), diabetic mitochondrial metabolism with control ultrastructure (Set CD – column 2) and diabetic mitochondrial metabolism with diabetic ultrastructure (Set DD – column 3). The regions belonging to mitochondria, glycogen and T-tubule are shown in black as part of the background. Each row of the figure represents a particular metabolic parameter: **A**. Distributions of myofibrillar ADP/ATP ratios. **B**. Distributions of myofibrillar Pi. **C**. Distributions of myofibrillar V_ATPase_. **D**. Distributions of myofibrillar V_CK_ / V_ATPase_ ratio.

The ratio of ADP and ATP concentration is considered as a key regulator of sarcomere shortening velocity and force development (48, 49). We can observe in Fig. 3A that myofibril areas located away from mitochondria have a higher ADP/ATP ratio compared to the cell wide average. Compared to simulation CC, this effect is more prominent in simulation CD and DD. We further observe variation in the myofibrillar concentration of Pi in all simulations (Fig. 3B). But gradients of Pi concentration appear to be weaker than ADP/ATP gradients. Myofibrillar ADP/ATP ratio can change drastically within 1 μm, depending on proximity to mitochondria. In contrast, Pi concentration does not exhibit strong localized gradients and rather changes gradually from one end of a cross section to the other. The simulations also predict a large difference in spatially averaged Pi concentrations in the three simulations. Average Pi is lowest in the simulation CC, and it cumulatively increases as mitochondrial dysfunction and diabetic ultrastructure are introduced in simulation CD and DD.

Fig. 3C shows the profiles of myofibrillar V_ATPase_ in the three simulations. The average V_ATPase_ is highest in the control simulation, and it declines with introduction of both mitochondrial dysfunction and diabetic ultrastructure. The model parameters influencing these results and their statistical significance has been discussed in detail in the next sub-section (Fig. 4). Due to our steady state assumptions, total ATP hydrolysis in the myofibrils (V_ATPase_*Area_myofibrils_) balances mitochondrial ATP synthesis (V_F1-F0_*Area_mitochondria_) in all the three simulations. This is evident in Fig. S3A, where average V_F1-F0_ drops to a lower level with introduction of mitochondrial dysfunction in simulation CD. While the average V_F1-F0_ appears to be same between simulation CD and DD, the total ATP synthesis is further diminished in DD due to a lower mitochondrial area fraction.

**Figure 4:**
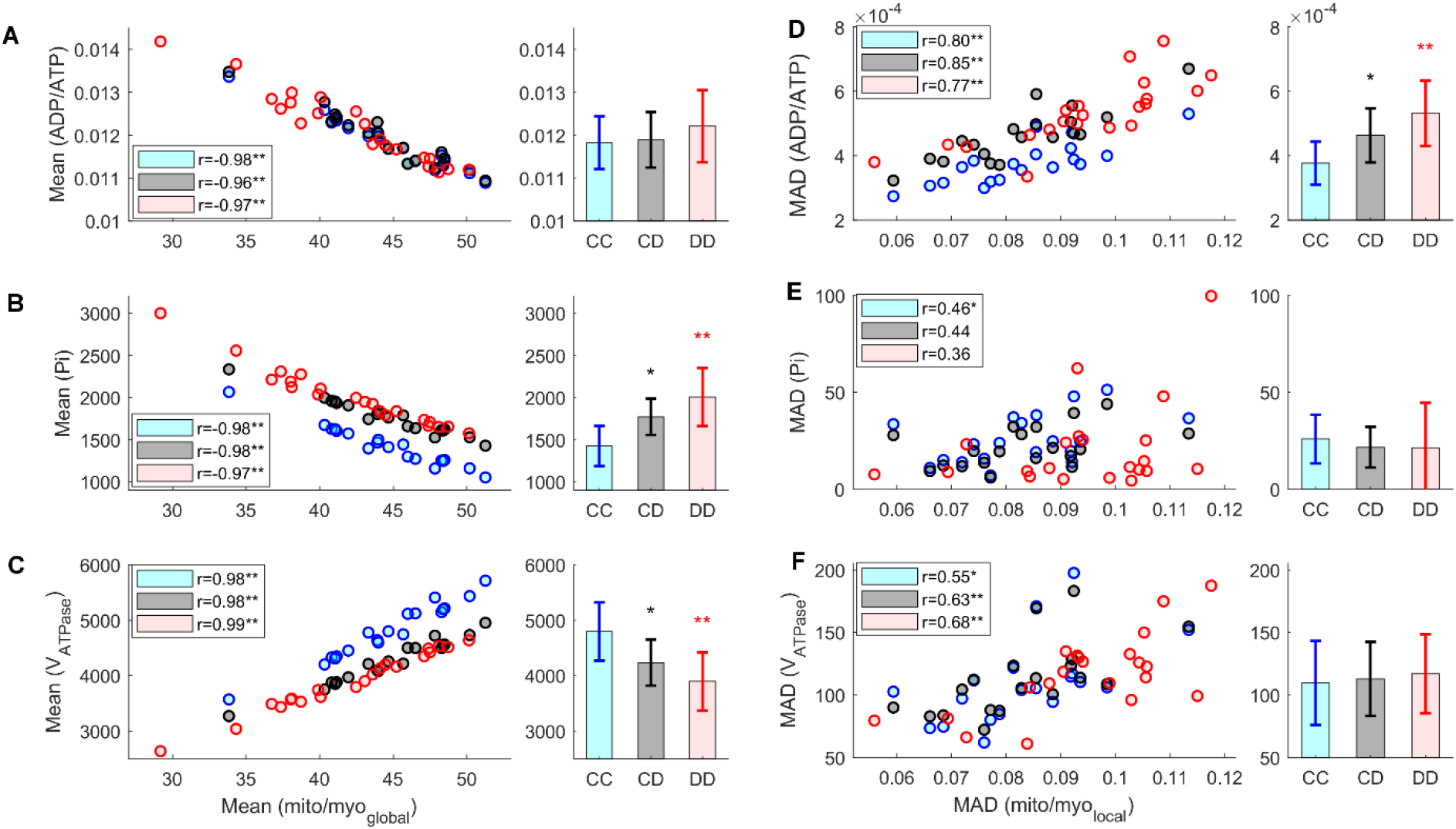
Relationship between cardiac ultrastructure and energy metabolism. The scatterplots show the spatial average and MAD of metabolic parameters in each cross section as a function of either corresponding average mito/myo_global_ ratio or MAD of mito/myo_local_ ratio. Each scatter plot contains results from simulation set CC (19 control cross sections), CD (19 control cross sections) and DD (21 diabetic cross sections). The legends show the Pearson correlation coefficient (r) separately for each simulation set. The p-value corresponding to the null hypothesis of no correlation is indicated as either * (0.05 ≥ p > 0.01) or ** (0.01 ≥ p). The bar plots located next to the scatterplots show the spatially averaged ADP/ATP ratios further averaged over three animals corresponding to each simulation set. The p-values indicated as *(0.05 ≥ p > 0.01) or **(0.01 ≥ p) correspond to the null hypothesis of unchanged value from CC. **A**. (**left**) ADP/ATP ratios in each cross section and (**right**) their simulation averages. **B**. (**left**) Average Pi concentration (μM) in each cross section and (**right**) their simulation averages. **C**. (**left**) Average V_ATPase_ (μmol/L. myofibrils/sec) in each cross section and (**right**) their simulation averages. **D**. (**left**) MAD of ADP/ATP ratios in each cross section and (**right**) their simulation averages. **E**. (**left**) MAD of Pi concentration in each cross section and (**right**) their simulation averages. **F**. (**left**) MAD of V_ATPase_ in each cross section and (**right**) their simulation averages.

### Type I diabetes alters the phosphocreatine shuttle

Due to cytosolic diffusion barriers, ATP produced in the mitochondria diffuses slowly towards the myofibrils (50). This diffusion rate (diffusivity = 30 μm/s) is usually not sufficient to fulfil the demand of ATP during a high cross bridge workload. However, myofibrillar CK enzyme can locally regenerate ATP by transferring the phosphate group of PCr to ADP (ADP + PCr ⇌ ATP + Cr). The generated Cr rapidly diffuses (diffusivity = 260 μm/s) to the mitochondrial IMS, where mtCK converts it back to PCr (ADP + PCr ⇌ ATP + Cr) through the reverse reaction (Fig. 2). The PCr synthesized by mitochondrial CK diffuses back to the myofibrils, thus completing a circular reaction pathway. This CK mediated pathway of ADP-ATP exchange is also known as the phosphocreatine shuttle. The ratio between reaction rates of myofibrillar CK enzyme and ATP hydrolysis (V_CK_ / V_ATPase_) indicates the fraction of ATP utilized by cross bridge cycle that is exchanged through this pathway.

Fig. 3D reveals that myofibril units adjacent to mitochondria have a lower V_CK_/V_ATPase_ compared to myofibrils located away from the mitochondrial columns. This trend is noticeable in all the three simulations. The results signify that a higher fraction of ATP is regenerated through myofibrillar CK, when diffusion distance between mitochondria and myofibrils is longer. On the other hand, direct exchange of ATP and ADP through diffusion takes a shorter time when myofibrils are in the vicinity of mitochondria. Therefore, less ADP is available to drive the myofibrillar CK enzymatic reactions forward.

In all steady state simulations, total PCr synthesis in the mitochondria (V_mtCK_*Area_myofibrils_) is balanced by the total PCr consumption (V_CK_ *Area_mitochondria_) in myofibrils. In simulations CD and DD, activity of mtCK is decreased by half. Consequently, CK reaction rates decline substantially in both mitochondria and myofibrils. This result is reflected in the visibly lower average V_CK_ / V_ATPase_ in simulation CD and DD compared to CC (Fig. 3D). The results imply that mitochondrial dysfunction weaken the phosphocreatine shuttle and encourage more direct diffusion based exchange of ADP and ATP. In addition, lower CK reaction rate leads to a large decline in myofibrillar PCr/ATP ratio as shown in Fig. S3B.

### Relationship between organelle area fractions and energy metabolism

Left hand panels of Fig. 4A and B show the spatially averaged ADP/ATP ratios and Pi concentrations corresponding to the 19 control and 21 diabetic cross sections. These have been plotted in conjunction with the ratio of mitochondrial area fraction against total mitochondrial and myofibrillar area fraction (mito/myo_global_) of each section. Averages of both ADP/ATP ratio and Pi concentration show a strong negative correlation with the mito/myo_global_ ratios. Since V_ATPase_ is modelled as an inverse function of ADP/ATP ratio and Pi concentration, average V_ATPase_ correlates positively with the mito/myo_global_ ratio of each cell cross section (Fig. 4C). The boxplots in the right hand panel of Fig. 4A reveal the absence of any significant change in spatially averaged ADP/ATP ratios with introduction of mitochondrial dysfunction (CD) and structural changes (DD). In contrast, Pi concentration is lowest in simulation set CC, and it successively rises in CD (24%) and DD (40%). We also observe a reverse trend with V_ATPase_ in Fig. 4C.

### Relationship between heterogeneity in ultrastructure and energy metabolism

MADs of the spatial distribution of myofibrillar ADP/ATP ratio, Pi concentration and V_ATPase_ act as indicators of the heterogeneity of the metabolic landscape in a cell cross section. Similarly, MAD of mito/myo_local_ distribution provides an indirect estimate of the structural heterogeneity with in a given cross section. As evident from the scatterplot in Fig. 4D, MAD of ADP/ATP appears to increase with higher MAD of mito/myo_local_ distribution in all three simulations sets. This positive correlation is evident in all three simulation sets. In contrast, no statistically significant correlation was observed between MAD of mito/myo_local_ and MADs of Pi in any of the three simulation sets (Fig. 4E).

The boxplots in the right hand panel of Fig. 4D reveal that structural alterations and mitochondrial dysfunction in simulation set DD lead to a 41% higher average MAD of ADP/ATP distribution compared to set CC. When results between simulation set CC and CD are compared, we can also observe a 23% rise in MAD of ADP/ATP ratio due to the introduction of mitochondrial dysfunction independent of structural alterations. The results suggest that when ATP demand is high, diabetic cell cross sections will have stronger concentration gradients of ADP and ATP compared to their control counterparts. Unlike ADP/ATP ratio, the MADs of Pi concentration and V_ATPase_ exhibited no statistically significant difference between results from the three simulation sets. These results are consistent with the weak Pi concertationg radients previously observed in Fig. 3B.

## Discussion

### Effects of diabetic mitochondrial dysfunction on cardiac energy metabolism

In simulation CD, the model of mitochondrial metabolism incorporates lower enzymatic activity of Complex I and F1-F0 ATP synthase along with elevated proton leak activity (Table 1). All of these changes diminish the capacity of mitochondria to synthesize ATP, which ultimately leads to a 12% decrease in both mitochondrial V_F1-F0_ and myofibrillar V_ATPase_. In myofibrils, the lower V_ATPase_ is driven by a 24% higher concentration of Pi. However, the average myofibrillar ADP/ATP remains at nearly same level. This result can be explained by the well-established role of PCr as a buffer of ATP (51, 52). The myofibrillar CK enzymes convert the excess build-up of ADP to ATP at the cost of the depleted PCr reserve (Fig. S3B). A substantial part of the ADP produced from ATP hydrolysis is also relocated to mitochondrial matrix, where it helps to drive the ATP synthesis forward despite of the lower F1-F0 activity.

Another important prediction of our model is decrease in CK mediated ADP-ATP exchange in T1D cardiomyopathy. Phospho-transfer between cardiac mitochondria and myofibrils occurs through three competitive pathways (Fig. S3). These include direct diffusion of ADP and ATP, alongside CK mediated phosphocreatine shuttle. According to an experimental study by Dzeja et al. on isolated mice hearts (53), phosphocreatine shuttle accounts for 69% of the total ADP-ATP exchange between mitochondria and myofibrils. Our control simulations (CC) also predict the average contribution of this shuttle (V_CK_/V_ATPase_) to be in a similar percentage range (Fig. 3D). However, in simulation CD, the contribution of phosphocreatine shuttle is reduced. Consequently, diffusion of ADP and ATP accounts for a higher proportion of total phospho-transfer in T1D cardiomyopathy.

Myofibrillar diffusivity of ADP and ATP is almost 10 times lower than that of Pi, PCr and Cr (50). Due to the low diffusivity, ADP and ATP require stronger concentration gradients than PCr and Cr to facilitate the same amount of phospho-transfer. As a result, myofibril areas with higher mito/myo_local_ density appear to have lower ADP/ATP, while myofibrils not in vicinity of mitochondria have high ADP/ATP ratio (Fig. 3A). This also explains why ADP/ATP ratio exhibits heterogenous spatial distribution unlike that of Pi (Fig. 3B). When it comes to simulation CD, the concentration gradients of ADP and ATP need to be even sharper to ensure their higher share of total phospho-transfer. Consequently, ADP/ATP distribution is significantly more heterogenous in simulation CD compared to CC (Fig. 4D).

### Ultrastructural alterations and their effect on cardiac energy metabolism

The analysis of TEM images in Fig. 1C reveal that diabetic cardiomyocyte cross sections have a significantly lower area fraction of mitochondria (37%) compared to their control counterparts (43%). At the same time, area fraction of myofibrils does not change significantly, implying a lower ATP synthesis capacity per unit area of myofibrils in diabetic cardiomyocytes. These results are in close agreement with previous imaging study by Searls et al. (24) on STZ induced T1D SD rat cardiomyocytes. Searls el al. found a significant reduction in mitochondrial area fraction (44.5% in control vs 36% in T1D) without any significant change in myofibrillar area fraction. Similar results have been also reported by Li et al (25).

As evident from the model predictions in Fig. 4C, the lower availability of mitochondria in diabetic cross-sections decrease the average V_ATPase_ in myofibrils (simulation DD). However, the average ADP/ATP ratio remains unchanged due to buffering by myofibrillar PCr. Previously, based on data from type I diabetic OVE26 mice, Shen et al. (14) proposed higher mitochondrial fraction as a compensatory response to mitochondrial dysfunction in T1D cardiomyopathy. Our model predictions contrast with this hypothesis and instead suggests lower mitochondrial area fraction as a mechanism that further exacerbates the negative impact of impaired OXPHOS (simulation DD vs CD).

Besides lower mitochondrial fraction, the arrangement of mitochondria and myofibrils is more non-uniform in T1D cardiomyopathy. This result is evident from the higher MAD of mito/myo_local_ ratio in the diabetic cross-sections (Fig. S1D). The increased spatial heterogeneity of mito/myo_local_ ratio is also reflected in the higher MAD of ADP/ATP ratio in simulation DD (Fig. 4D). In simulation CD, MAD of ADP/ATP ratio increases due to elevation in diffusive fluxes of ADP and ATP. The non-uniform arrangement of mitochondria and myofibrils further contributes to this heterogenous metabolic landscape in simulation DD. The observed spatial variation in ADP/ATP might lead to variation in actomyosin contraction velocity between different parts of a cell (48, 49). This may result in intracellular shear strain, along with negative consequences for the cellular ultrastructure. This model-informed hypothesis will require new precision experimental measurements.

### Limitations and further work

A logical next step of the current work will be to couple our model of cardiac bioenergetics with models of Ca2+ signalling and cross-bridge cycling (48, 49). This will enable us to translate the predictions of altered ADP/ATP level and Pi concentration in T1D cardiomyopathy to corresponding changes in cardiac force dynamics and contractility. Moreover, many previous studies report elevated intrinsic stiffness of cardiomyocytes due to diabetes (54, 55). A coupled bioenergetics-mechanics model of cardiomyocytes will be a useful tool to investigate the overall effects of the alterations in metabolism, ultrastructure and material properties. There is also scope to improve the current model of cardiac bioenergetics by incorporating details of glycolysis, beta oxidation and subsequent TCA cycle.

On the image analysis side, a more complete analysis of the 3D organization of mitochondrial columns and other organelles between control and T1D is needed. According to a recent work by Glancy et al. (10), cardiac mitochondria form a series of interconnected networks which can conduct electricity similar to power grids. These networks can dynamically respond to stress by electrically separating malfunctioning mitochondria. Another previous study indicates that the extent of T1D induced ultrastructural alterations can vary between different mitochondrial sub-populations within the same cell (19).

Regardless of the limitations discussed above, the EM image analysis in this study provides many new insights on cardiac ultrastructure that were not reported previously, e.g., more non-uniform distribution of mitochondria and myofibrils in T1D cardiomyopathy. The current study is also the first attempt in literature to create a spatially detailed mathematical model of cardiac energy metabolism in T1D cardiomyopathy. The model provides a simplistic representation of the diminished ATP and PCr synthesis capacity of diabetic mitochondria, which we used to understand the relationship between cardiac ultrastructure and bioenergetics.

## Conclusion

Alterations in the columnar ultrastructure of cardiac mitochondria is observed in several cardiac disease conditions. The primary aim of the current study was to understand the functional role of these ultrastructural alterations in the context of STZ induced T1D cardiomyopathy as a model disease state. Our bioenergetics simulations predict that diabetic cardiomyocytes have a higher level of myofibrillar Pi and lower ATP hydrolysis rate compared to control cardiomyocytes, in response to the same level of cross bridge cycle stimulation. These results are a consequence of lower availability of mitochondria per unit area of myofibrils, as well as impaired OXPHOS capacity of individual mitochondrion. Our simulations further reveal that spatial average of myofibrillar ADP/ATP ratio is not affected in T1D cardiomyopathy. However, the spatial distribution of ADP/ATP ratio becomes more heterogeneous in diabetic cross sections due to a more irregular arrangement of mitochondria and myofibrils and an increased dependence of the cell on direct ATP-ADP diffusion to meet the ATP demands of myofibrils. Overall, our study indicates that alterations in cardiac ultrastructure such as lower mitochondrial area fraction and irregular mitochondrial-myofibrillar arrangement further aggravate the metabolic disruptions preceding the ultrastructural alterations. Future work will include using a coupled bioenergetics-mechanics model to assess the impact of the observed metabolic disruptions on the cardiac force dynamics and contractility.

## Acknowledgments

F.S. was supported by a UKRI Future Leaders Fellowship, grant number [MR/T043571/1]. F.S. and V.R. are supported by a Royal Society International Exchange Award, grant number [IES\R3\203170]. This research was also supported by the Royal Society of New Zealand Marsden Fast Start Grant (11-UOA-184).

## Supplementary material

**Figure S1:**
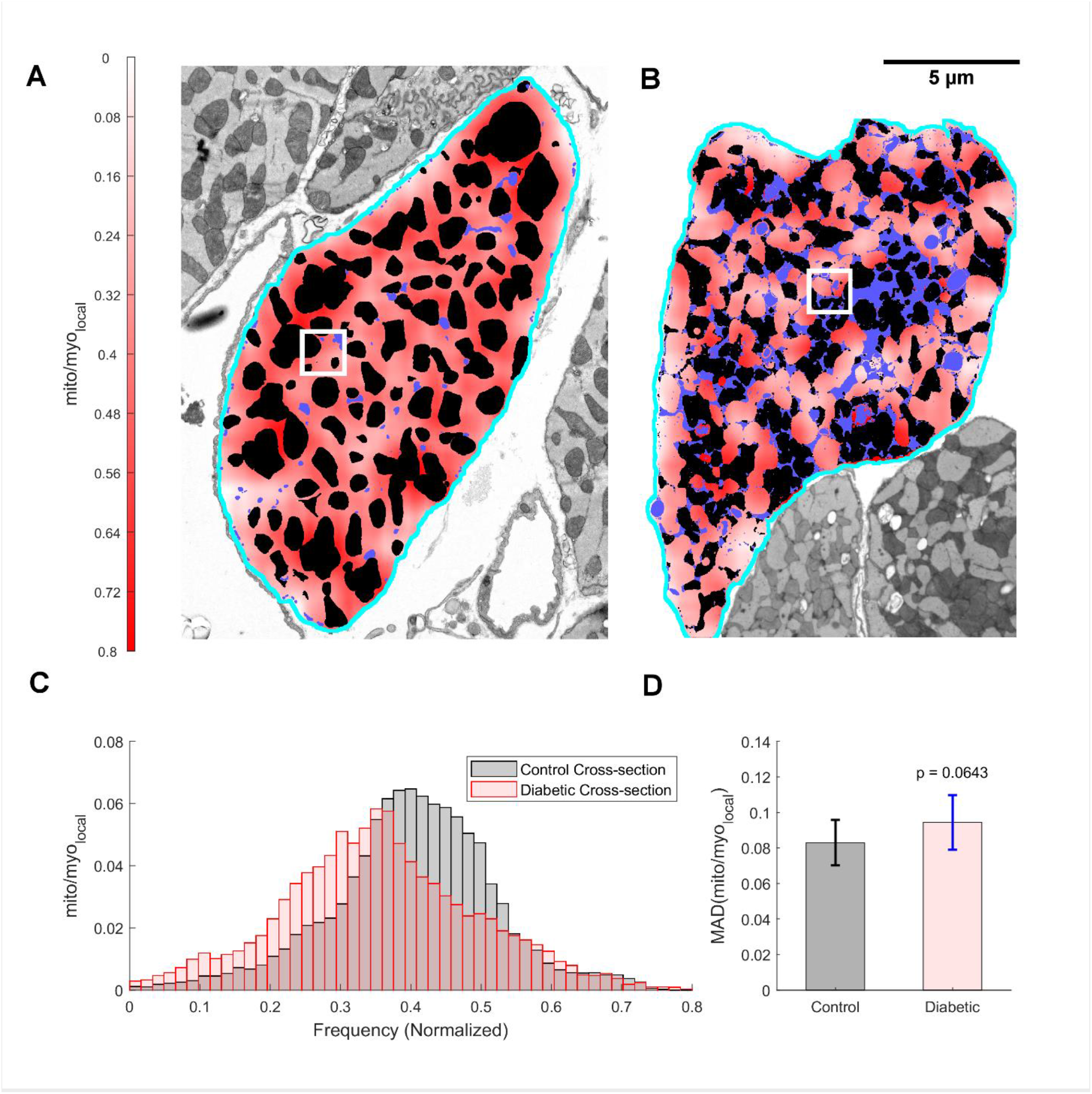
**A**. Distribution of mito/myo_local_ in myofibril regions of the representative control cardiomyocyte cross section. Mitochondria are marked in black, while all other organelles except myofibrils are shown in blue. The square window shows a sampling window of 1.6 μm used for calculating mito/myo_local_ in each myofibrillar pixel. **B**. Distribution of mito/myo_local_ in myofibril regions of the representative diabetic cardiomyocyte cross section. **C**. Histogram representation of mito/myo_local_ distributions in the representative control and diabetic cross sections. **D**. Average MAD of the mito/myo_local_ distributions in all control and diabetic cross sections. The diabetic MAD shows evidence of differing from the control case, with associated p = 0.064.

**Figure S2:**
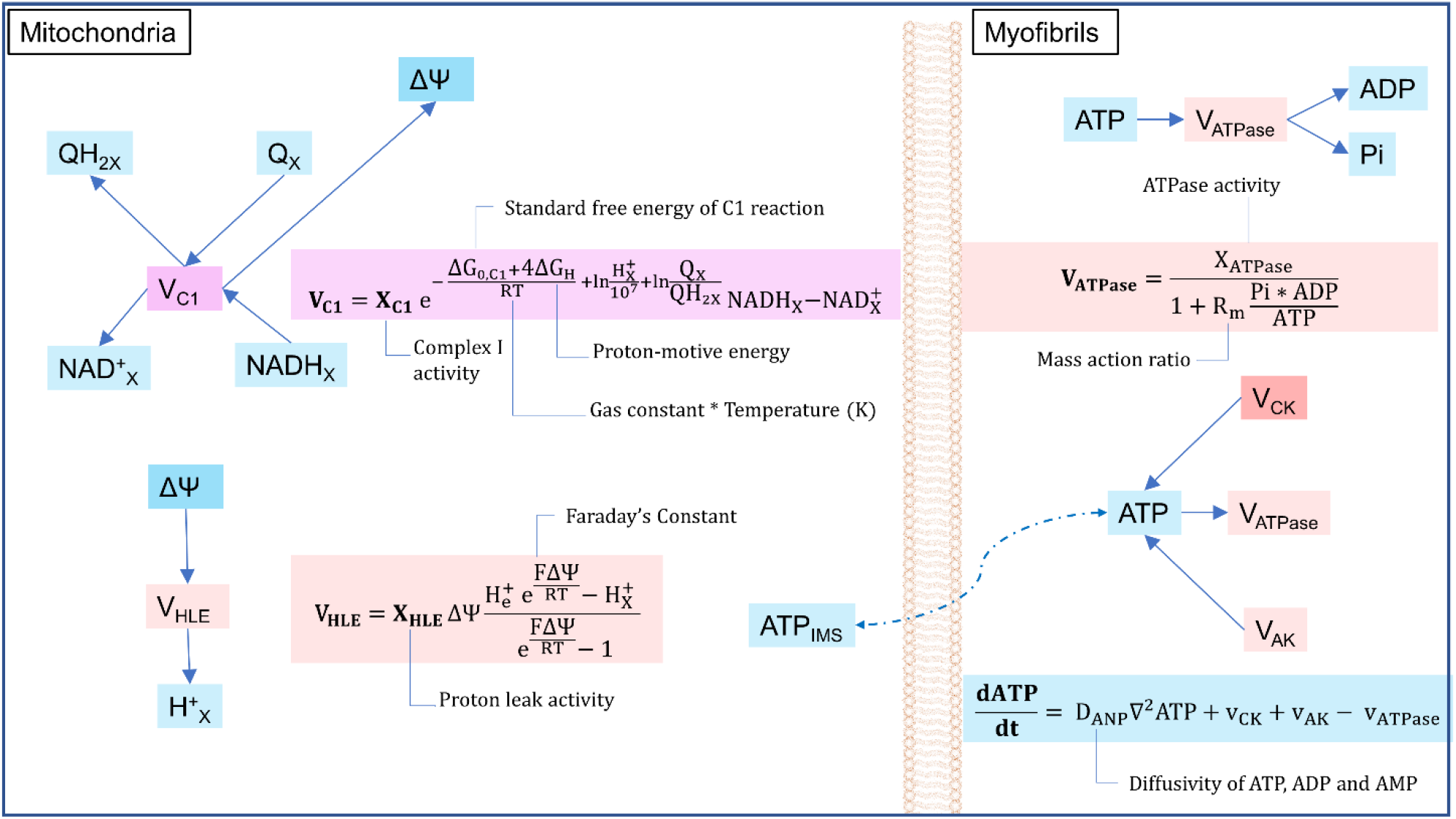
The figure shows the schematic diagram and formulation of biophysical equations of three reactions fluxes (V_C1_, V_HLE_ and V_ATPase_) and one state variable (myofibrillar ATP concentration) used in the cardiac bioenergetics model. The cyan boxes represent the molar concentration of metabolites in respective compartments.

**Figure S3:**
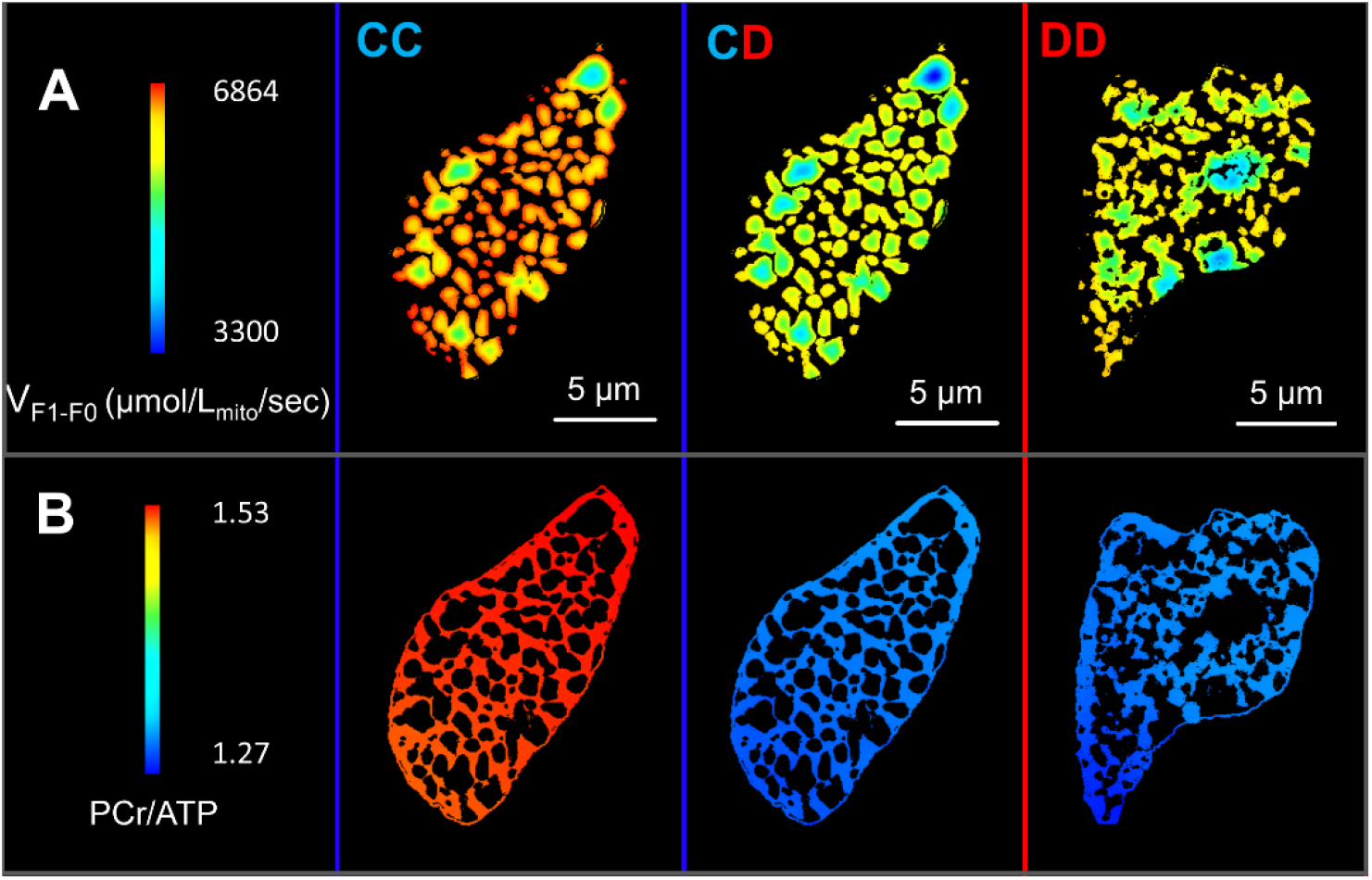
**A**. Model predicted distributions of mitochondrial V_F1-F0_ in three representative simulations. mitochondria, glycogen and T-tubule regions are shown in black as part of background **B**. Distributions of myofibrillar PCr/ATP ratio in the three simulations.

**Table S1:** List of state variables and reaction fluxes used in the model of energy metabolism

| Symbol | Compartment | Definition |
| --- | --- | --- |
| State Variables of the system |  |  |
| ATP | Myofibrils | Adenosine triphosphate (μM) |
| ATP<sub>IMS</sub> | Mitochondria IMS |  |
| ATP<sub>X</sub> | Mitochondria matrix |  |
| ADP | Myofibrils | Adenosine diphosphate (μM) |
| ADP<sub>IMS</sub> | Mitochondria IMS |  |
| ADP<sub>X</sub> | Mitochondria matrix |  |
| AMP | Myofibrils | Adenosine monophosphate (μM) |
| AMP<sub>IMS</sub> | Mitochondria IMS |  |
| PCr | Myofibrils | Phosphocreatine (μM) |
| PCr<sub>IMS</sub> | Mitochondria IMS |  |
| Cr | Myofibrils | Creatine (μM) |
| Cr<sub>IMS</sub> | Mitochondria IMS |  |
| Pi | Myofibrils | Inorganic phosphate (μM) |
| Pi<sub>IMS</sub> | Mitochondria IMS |  |
| Pi<sub>X</sub> | Mitochondria matrix |  |
| H<sup>+</sup><sub>X</sub> |  | H<sup>+</sup> concentration (μM) |
| NADH<sub>X</sub> |  | Nicotinamide adenine dinucleotide reduced (μM) |
| NAD<sup>+</sup><sub>X</sub> |  | Nicotinamide adenine dinucleotide oxidized (μM) |
| Q<sub>X</sub> |  | Ubiquinone (μM) |
| QH2<sub>X</sub> |  | Ubiquinol (μM) |
| cytC<sup>2+</sup><sub>X</sub> |  | Cytochrome C reduced (μM) |
| cytC<sup>3+</sup><sub>X</sub> |  | Cytochrome C oxidized (μM) |
| ΔΨ | Mitochondrial inner membrane | Mitochondrial inner membrane (mV) |
| Fluxes of reactions |  |  |
| V<sub>ATPase</sub> | Myofibrils<br>(mol/sec/Litre of myofibril) | Flux of ATP consumption |
| V<sub>CK</sub> |  | Flux of creatine kinase reaction |
| V<sub>AK</sub> |  | Flux of adenylate kinase reaction |
| V<sub>mtCK</sub> | Mitochondrial IMS<br>(mol/sec/Litre of mitochondria) | Flux of mitochondrial creatine kinase reaction |
| V<sub>mtAK</sub> |  | Flux of mitochondrial adenylate kinase reaction |
| V<sub>DH</sub> | Mitochondrial matrix<br>(mol/sec/Litre of mitochondria) | Dehydrogenase flux representing the TCA cycle and other NADH-producing reactions |
| V<sub>F1FO</sub> | Mitochondrial IMS<br>+<br>matrix<br>(mol/sec/Litre of mitochondria) | Flux through complex V (F1F0 - ATP synthase) |
| V<sub>C4</sub> |  | Flux through complex IV |
| V<sub>C3</sub> |  | Flux through complex III |
| V<sub>C1</sub> |  | Flux through complex I |
| V<sub>PiH</sub> |  | Flux through Phosphate hydrogen co-transporter |
| V<sub>ANT</sub> |  | Flux through Adenine nucleotide translocases |

**Table S2:** List of diffusion constants and their values

| Diffusion Constant | Metabolites | Myofibrillar value | Mitochondrial value |
| --- | --- | --- | --- |
| $D_{\text{ANP}}$ | ATP, ADP and AMP | $30 \mu\text{m}^2/\text{s}$ | $3 \mu\text{m}^2/\text{s}$ |
| $D_{\text{PCr}}$ | PCr | $260 \mu\text{m}^2/\text{s}$ | $26 \mu\text{m}^2/\text{s}$ |
| $D_{\text{Cr}}$ | Cr | $260 \mu\text{m}^2/\text{s}$ | $26 \mu\text{m}^2/\text{s}$ |
| $D_{\text{Pi}}$ | Pi | $327 \mu\text{m}^2/\text{s}$ | $32.7 \mu\text{m}^2/\text{s}$ |

