## Supplementary material for "Effects of altered cellular ultrastructure on energy metabolism in diabetic cardiomyopathy – an in-silico study": Methods S1

### SUPPLEMENTAL METHODS

#### Statistical Analyses

This report includes details of the statistical analyses conducted in the main article. In this study p-values were calculated for statistical comparisons but the statistical significance was not determined based on an arbitrary significance threshold; instead, p-values were interpreted as continuous variables; for each hypothesis tested the minimum False Positive Risk (mFPR) was calculated to assist the interpretation of the statistical test (1-2). In what follows, the first section introduces the two sets of data generated to investigate how cellular architecture and metabolism are related in diabetic cardiomyopathy. The second section presents the hypothesis and results of the correlation analyses (3). The third section presents how Linear Mixed Models (4-5) were applied for inferential purposes on the data. All the statistical analyses have been performed in R version 4.1.0 (5).

#### Data

This study employs two different sets of data. The first dataset contains measurements estimated from cross-sectional electron microscopic (EM) images collected from rat cardiac tissue corresponding to two different conditions: control and diabetes. There were 3 biological replicates for each condition of rats. For each biological replicate, different cross-sections of the cardiac tissue were acquired (technical replicates) using electron microscopy (EM). The summary of the collected images can be found in Table 1.

| Control Sample | n. cross-sections | Diabetes sample | n. cross-sections |
|---|---|---|---|
| 1 | 5 | s 1 | 8 |
| 2 | 8 | s 2 | 8 |
| 3 | 6 | s 3 | 5 |

**Table 1**: Images collected through EM. Control Sample include three animals employed for the experiment in control group, while Diabetes Sample indicates the animals with induced type I Diabetes.

The second dataset contains predictions from three different set of in-silico experiments – CC (Control mitochondrial metabolism with Control ultrastructure), CD (Diabetic mitochondrial metabolism with Control ultrastructure) and DD (Diabetic mitochondrial metabolism with Diabetic ultrastructure). Simulation CC and CD were based on cross-sectional EM images collected from control animals, while DD was based on diabetic EM cross-sections. The samples generated from the three in-silico simulation sets are summarised in Table 2

| CC | n. cross-sections | CD | n. cross-sections | DD | n. cross-sections |
|---|---|---|---|---|---|
| CC 1 | 5 | CD 1 | 5 | DD 1 | 8 |
| CC 2 | 8 | CD 2 | 8 | DD 2 | 8 |
| CC 3 | 6 | CD 3 | 6 | DD 3 | 5 |

**Table 2**: Samples collected through in-silico experiments. CC denotes control samples, CD indicates the sample with control cellular architecture, but mitochondrial dysfunction, while DD includes type I diabetic cross-section with mitochondrial dysfunctions.

#### Correlation Analysis

The correlation between any two measurements were calculated using the Pearson Correlation Coefficient. Subsequently, the two-tailed hypothesis that the correlation is null was tested employing the t-test. The correlation test considers each observation based on individual EM cross-sections as an independent observation. The following list systematically summarises all the correlation analyses performed, while Table 3 reports their results.

1. Within each condition, namely CC, CD and DD, is the correlation between "mito/myo$_{global}$" and "MEAN ADP/ATP" null?
2. Within each condition, namely CC, CD and DD, is the correlation between "mito/myo$_{global}$" and "MEAN Pi" null?
3. Within each condition, namely CC, CD and DD, is the correlation between "mito/myo$_{global}$" and "MEAN $V_{ATPase}$" null?
4. Within each condition, namely CC, CD and DD, is the correlation between "MAD mito/myo$_{local}$" and "MAD ADP/ATP" null?
5. Within each condition, namely CC, CD and DD, is the correlation between "MAD mito/myo$_{local}$" and "MAD Pi" null?
6. Within each condition, namely CC, CD and DD, is the correlation between "MAD mito/myo$_{local}$" and "MAD $V_{ATPase}$" null?

| Question | Pivot | Corr | Std. Err. | $CI_{0.025}$ | $I_{0.975}$ | $T_{obs}$ | $p_{value}$ | mFPR |
|---|---|---|---|---|---|---|---|---|
| 1 cc | $T_{17}$ | 0.97800 | 0.05058 | 1.08473 | 87128 | .33384 | $5\ 10^{-13}$ | $3\ 10^{-12}$ |
| 1 cd | $T_{17}$ | 0.96405 | 0.06443 | 1.10001 | 82810 | .96073 | $3\ 10^{-11}$ | $2\ 10^{-10}$ |
| 1 dd | $T_{19}$ | 0.96699 | 0.05845 | 1.08934 | 84463 | .54158 | $9\ 10^{-13}$ | $7\ 10^{-12}$ |
| 2 cc | $T_{17}$ | 0.98086 | 0.04722 | 1.08049 | 88123 | .77189 | $1\ 10^{-13}$ | $1\ 10^{-12}$ |
| 2 cd | $T_{17}$ | 0.98102 | 0.04702 | 1.08023 | 88181 | .86223 | $1\ 10^{-13}$ | $1\ 10^{-12}$ |
| 2 dd | $T_{19}$ | 0.97469 | 0.05128 | 1.08202 | 86736 | .00734 | $8\ 10^{-14}$ | $6\ 10^{-13}$ |

| | | | | | | | | |
|---|---|---|---|---|---|---|---|---|
| 3 cc | $T_{17}$ | ).97977 | 0.04853 | ).87737 | )8217 | 18690 | $2\ 10^{-13}$ | $1\ 10^{-12}$ |
| 3 cd | $T_{17}$ | ).97933 | 0.04905 | ).87584 | )8282 | 96522 | $3\ 10^{-13}$ | $2\ 10^{-12}$ |
| 3 dd | $T_{19}$ | ).99155 | 0.02975 | ).92927 | )5383 | 32394 | $2\ 10^{-18}$ | $1\ 10^{-17}$ |
| 4 cc | $T_{17}$ | ).79959 | 0.14565 | ).49229 | 10689 | 18970 | $3\ 10^{-5}$ | ).00028 |
| 4 cd | $T_{17}$ | ).84599 | 0.12931 | ).57315 | 11883 | 54186 | $5\ 10^{-06}$ | $3\ 10^{-15}$ |
| 4 dd | $T_{19}$ | ).77469 | 0.14506 | ).47107 | )7832 | 34031 | $3\ 10^{-05}$ | ).00027 |
| 5 cc | $T_{17}$ | ).45626 | 0.21581 | ).00092 | )1160 | 11411 | ).04958 | ).20646 |
| 5 cd | $T_{17}$ | ).43902 | 0.21791 | 0.02072 | 39878 | )1471 | 0.0600 | ).23541 |
| 5 dd | $T_{19}$ | ).35606 | 0.21438 | 0.09264 | 30476 | 56088 | ).11314 | ).34802 |
| 6 cc | $T_{17}$ | ).54939 | 0.20265 | ).12182 | )7695 | 71098 | ).01483 | ).07991 |
| 6 cd | $T_{17}$ | ).62819 | 0.18870 | ).23005 | )2632 | 32894 | ).00397 | ).02450 |
| 6 dd | $T_{19}$ | ).68386 | 0.16738 | ).33353 | )3420 | )8565 | ).00063 | ).00435 |

**Table 3**: Results of the correlation analysis across different variables.

#### Linear Mixed Model Analysis

Given the nested structure of the datasets, one of the best options to evaluate the marginal effect of a condition, such as Control and Diabetes or CC, CD, DD, is to employ the linear mixed models. In linear mixed models, the random effect is attributed to the repeated measures (cross-sections) of the same sample of rat cardiac tissue, while the fixed effect is the condition of the sample, where Control or the CC is in general set as a baseline. The pairwise difference in average between two conditions was evaluated using a t-test on the coefficient associated with the condition of interest. The question and associated results of the analyses are listed below.

**a) Does Mitochondria % show any difference between control and diabetic group?**

| Effect | Variable | Parameter | Std. Err. | df | $t_{obs}$ | $p_{value}$ | mFPR |
|---|---|---|---|---|---|---|---|
| Random | Cross-Section | *1.57319* *(stdDev intercept)* | *4.33176* *(stdDev residuals)* | | | | |
| Fixed | Intercept | 43.22972 | 1.35268 | 34 | 31.95836 | 0 | |
| Fixed | Diabetes | -6.14261 | 1.88877 | 4 | -3.25216 | 0.0313 | 0.10604 |

**b) Does Myofibrils % show any difference between control and diabetic groups?**

| Effect | Variable | Parameter | Std. Err. | df | $t_{obs}$ | $p_{value}$ | mFPR |
|---|---|---|---|---|---|---|---|
| Random | Cross-Section | *2.60491* *(stdDev intercept)* | *4.33537* *(stdDev residuals)* | | | | |
| Fixed | Intercept | 53.19633 | 1.81033 | 34 | 29.38486 | 0 | |
| Fixed | Diabetes | -1.86826 | 2.54250 | 4 | -0.73481 | 0.5032 | 0.60071 |

**c) Does Glycogen % show any difference between control and diabetic groups?**

| Effect | Variable | Parameter | Std. Err. | df | $t_{obs}$ | $p_{value}$ | mFPR |
|---|---|---|---|---|---|---|---|
| Random | Cross-Section | *2.73750* *(stdDev intercept)* | *2.29215* *(stdDev residuals)* | | | | |
| Fixed | Intercept | 1.78526 | 1.66852 | 34 | 1.06996 | 0.2922 | |
| Fixed | Diabetes | 6.68926 | 2.35441 | 4 | 2.84115 | 0.0468 | 0.15046 |

**d) Does T-tubules % show any difference between control and diabetic groups?**

| Effect | Variable | Parameter | Std. Err. | df | $t_{obs}$ | $p_{value}$ | mFPR |
|---|---|---|---|---|---|---|---|
| Random | Cross-Section | *0.28447* *(stdDev intercept)* | *0.88986* *(stdDev residuals)* | | | | |
| Fixed | Intercept | 1.69601 | 0.26321 | 34 | 6.44358 | 0 | |
| Fixed | Diabetes | 1.34198 | 0.36695 | 4 | 3.65706 | 0.0216 | 0.07571 |

**e) Does mito/myo$_{local}$ MAD show any difference between control and diabetic groups?**

| Effect | Variable | Parameter | Std. Err. | df | $t_{obs}$ | $p_{value}$ | mFPR |
|---|---|---|---|---|---|---|---|
| Random | Cross-Section | *$3\ 10^{-7}$* *(stdDev intercept)* | *0.01415* *(stdDev residuals)* | | | | |
| Fixed | Intercept | 0.08301 | 0.00324 | 34 | 25.55811 | 0 | |
| Fixed | Diabetes | 0.01136 | 0.00448 | 4 | 2.53471 | 0.0643 | 0.19519 |

**f) Does MEAN ADP/ATP show any difference amongst CC (baseline), CD and DD?**

| Effect | Variable | Parameter | Std. Err. | df | $t_{obs}$ | $p_{value}$ | mFPR |
|---|---|---|---|---|---|---|---|
| Random | Cross-Section | *0.00031* *(stdDev intercept)* | *0.00065* *(stdDev residuals)* | | | | |
| Fixed | Intercept | 0.01178 | 0.00023 | 50 | 49.16090 | 0 | |
| Fixed | CD | 0.00006 | 0.00033 | 6 | 0.19579 | 0.8512 | 0.66204 |
| Fixed | DD | 0.00044 | 0.00033 | 6 | 1.33478 | 0.2304 | 0.46280 |

**g) Does MEAN Pi show any difference amongst CC (baseline), CD and DD?**

| Effect | Variable | Parameter | Std. Err. | df | $t_{obs}$ | $p_{value}$ | mFPR |
|---|---|---|---|---|---|---|---|
| Random | Cross-Section | *93.86631* *(stdDev intercept)* | *263.1411* *(stdDev residuals)* | | | | |
| Fixed | Intercept | 1416.1821 | 81.50814 | 50 | 17.37473 | 0 | |
| Fixed | CD | 346.2719 | 115.26992 | 6 | 3.00400 | 0.0239 | 0.09572 |
| Fixed | DD | 593.561 | 113.78595 | 6 | 5.21647 | 0.0020 | 0.00887 |

**h) Does MEAN of $V_{ATPase}$ show any difference amongst CC (baseline), CD and DD?**

| Effect | Variable | Parameter | Std. Err. | df | $t_{obs}$ | $p_{value}$ | mFPR |
|---|---|---|---|---|---|---|---|
| Random | Cross-Section | *218.7274* *(stdDev intercept)* | *457.0094* *(stdDev residuals)* | | | | |
| Fixed | Intercept | 4826.222 | 164.8878 | 50 | 29.26974 | 0 | |
| Fixed | CD | -566.214 | 233.1865 | 6 | -2.42816 | 0.0513 | 0.18135 |
| Fixed | DD | -942.579 | 231.0037 | 6 | -4.08036 | 0.0065 | 0.02845 |

**i) Does MAD of ADP/ATP show any difference amongst CC (baseline), CD and DD?**

| Effect | Variable | Parameter | Std. Err. | df | $t_{obs}$ | $p_{value}$ | mFPR |
|---|---|---|---|---|---|---|---|
| Random | Cross-Section | *$2\ 10^{8}$* *(stdDev intercept)* | *$8\ 10^{-5}$* *(stdDev residuals)* | | | | |
| Fixed | Intercept | 0.00037 | $1\ 10^{-5}$ | 50 | 19.06908 | 0 | |
| Fixed | CD | 0.00008 | $2\ 10^{-5}$ | 6 | 3.08347 | 0.0216 | 0.08743 |
| Fixed | DD | 0.00015 | $2\ 10^{-5}$ | 6 | 5.67146 | 0.0013 | 0.00580 |

**j) Does MAD of Pi show any difference amongst CC (baseline), CD and DD?**

| Effect | Variable | Parameter | Std. Err. | df | $t_{obs}$ | $p_{value}$ | mFPR |
|---|---|---|---|---|---|---|---|
| Random | Cross-Section | *3.684054* *(stdDev intercept)* | *16.36469* *(stdDev residuals)* | | | | |
| Fixed | Intercept | 25.714096 | 4.330213 | 50 | 5.938298 | 0 | |
| Fixed | CD | -4.161927 | 6.123846 | 6 | -0.679626 | 0.5221 | 0.611144 |
| Fixed | DD | -4.248331 | 6.014195 | 6 | -0.706384 | 0.5064 | 0.60672 |

**k) Does MAD of $V_{ATPase}$ show any difference amongst CC (baseline), CD and DD?**

| Effect | Variable | Parameter | Std. Err. | df | $t_{obs}$ | $p_{value}$ | mFPR |
|---|---|---|---|---|---|---|---|
| Random | Cross-Section | *0.00243* *(stdDev intercept)* | *31.64691* *(stdDev residuals)* | | | | |
| Fixed | Intercept | 109.48611 | 7.26029 | 50 | 15.08010 | 0 | |
| Fixed | CD | 3.39116 | 10.26761 | 6 | 0.33027 | 0.7524 | 0.65351 |
| Fixed | DD | 7.47337 | 10.02016 | 6 | 0.74583 | 0.4840 | 0.59990 |
